# An efficient and cost-effective method for directed mutagenesis at multiple dispersed sites – a case study with Omicron Spike DNA

**DOI:** 10.1101/2022.02.10.480017

**Authors:** Rita Rani, Kishore V.L. Parsa, Kiranam Chatti, Aarti Sevilimedu

## Abstract

Site directed mutagenesis is an invaluable technique which enables the elucidation of the contribution of specific residues to protein structure and function. The simultaneous introduction of mutations at a large number of sites (>10), singly and in multiple combinations is often necessary to fully understand the functional contributions. We report a simple, efficient, time and cost effective method to achieve this using commonly available molecular biology reagents and protocols, as an alternative to gene synthesis. We demonstrate this method using the Omicron Spike DNA construct as an example, and create a construct bearing 37 mutations (as compared to wild-type Spike DNA), as well as four other constructs bearing subsets of the full spectrum of mutations. We believe that this method can be an excellent alternative to gene synthesis, especially when three or more variants are required.

**Graphical Abstract:** 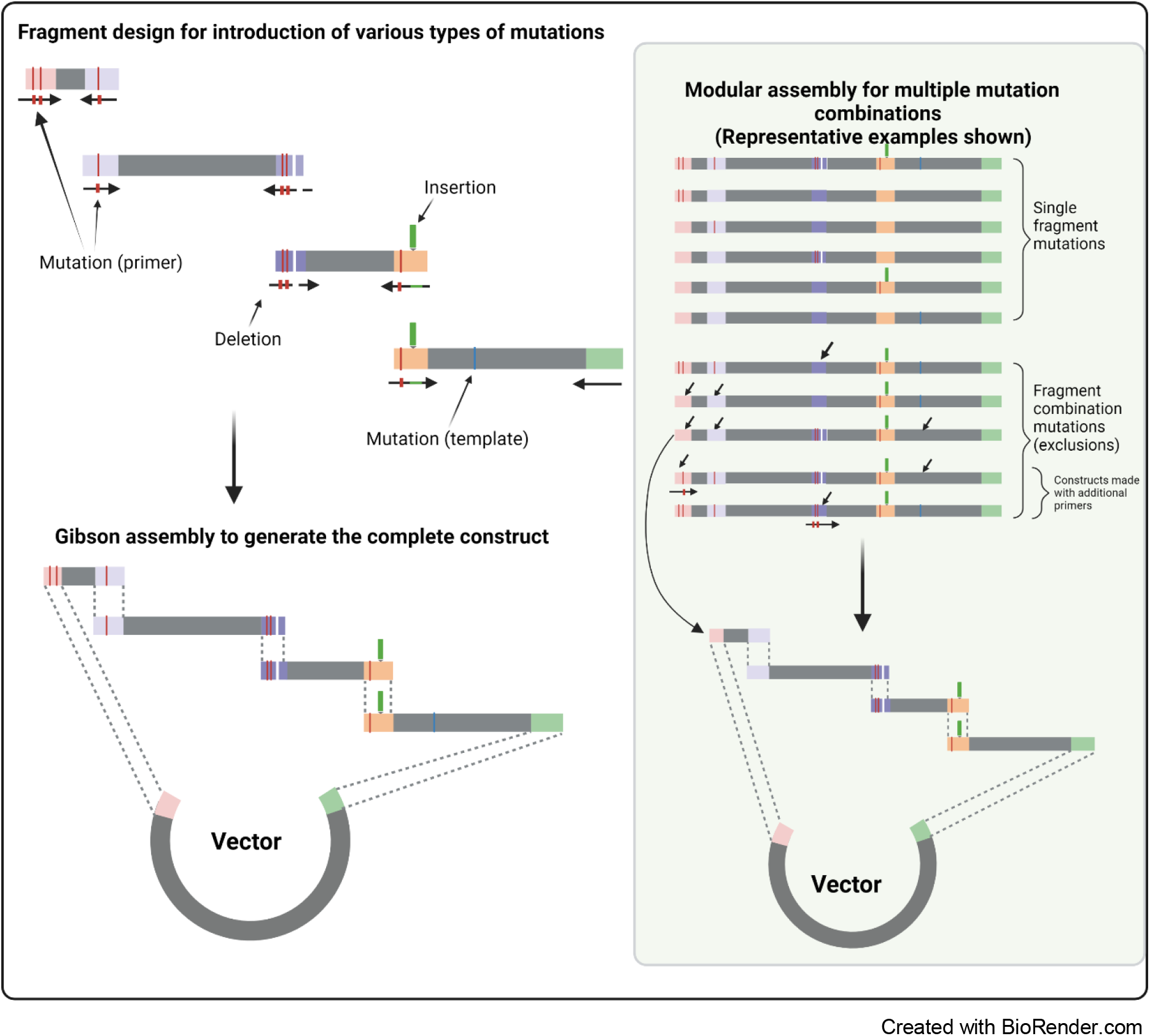

## Introduction

The method of choice for the introduction of one or a few mutations located in close proximity in recombinant DNA constructs, is site-directed mutagenesis. In contrast, for introducing a large number of mutations, especially those not clustered together within the region covered by a typical primer and spread over several kilobases, commercial gene synthesis is preferred [1]. However, this may be too expensive, may take too long, may not be an option in some geographic locations or for some projects. In this scenario, an alternative method combining PCR, overlap extension PCR and Gibson cloning can be used to create the desired construct in a short duration and at a cost lower than that of commercial synthesis. This alternate strategy will provide a significant cost advantage when a series of constructs with combinations of mutations needs to be created.

The basic principle of the method lies in assembling the complete construct using a series of smaller, mutation containing fragments. The construct is split into fragments in such a way that forward and reverse primers used to amplify a fragment will contain all of the mutations located in that fragment. The mutation containing fragments are then seamlessly assembled using a homology based cloning method such as Gibson assembly [2]. Mutations located within a 50bp region are introduced using a single primer; fragment sizes typically range from 100bp to 3kb and an unlimited number of fragments can be combined into the final construct in one or more rounds of cloning. Due to the modular nature of construct assembly, a combinatorial library of constructs including or excluding any combination of mutations can be generated with minimal additional effort, time and cost.

We illustrate this method using the example of Omicron Spike gene which contains 37 mutations (dispersed over nearly 3kb) as compared to the wild-type Spike gene. We followed a similar strategy to create the Delta Spike construct previously, which was successfully used in the pseudovirus neutralization assay reported by Kumar et al [3], validating this approach.

The quote obtained for commercial synthesis of this construct from a reputed vendor was INR 1 Lakh (USD 1330) and the time from order to delivery of the sequence verified construct was 45 days at our location, possibly due to import restrictions and pandemic related delays. The in-house synthesis of the final sequence verified construct was completed in under 29 days at an estimated cost of INR 50,000 (USD 665) (Table 3). In addition to the complete Omicron Spike gene, four additional variants were synthesized using no additional reagents. The time required for in-house synthesis was primarily limited by the time required for commercial oligo synthesis/delivery and turnaround time for sequencing, which required 50% of the total time to synthesize and sequence-validate the full-length construct. The cost saving was in part due to the use of lab-made enzymes and cloning reagents.

**Table 1:**
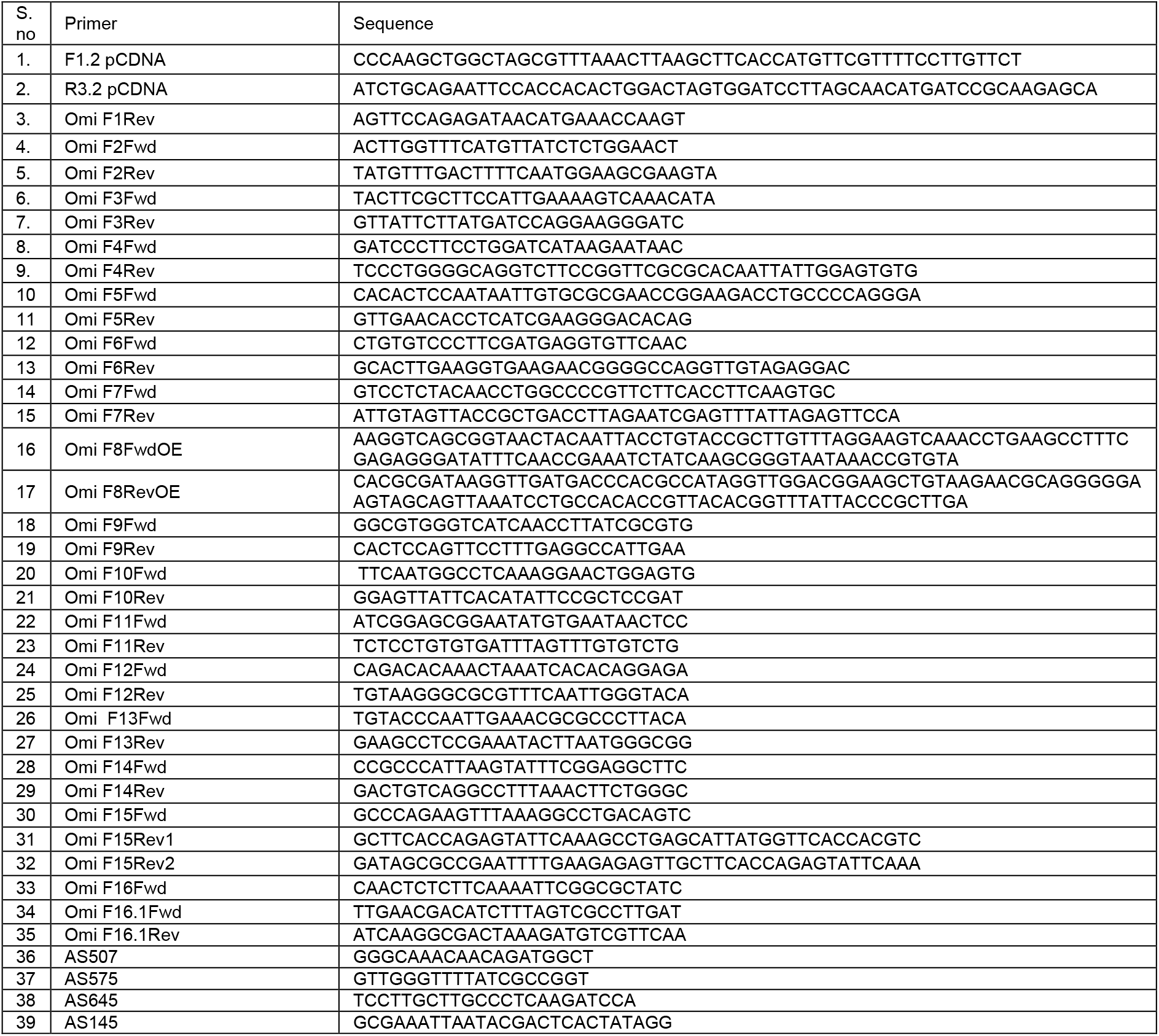
List of primers used in this study

**Table 2:**
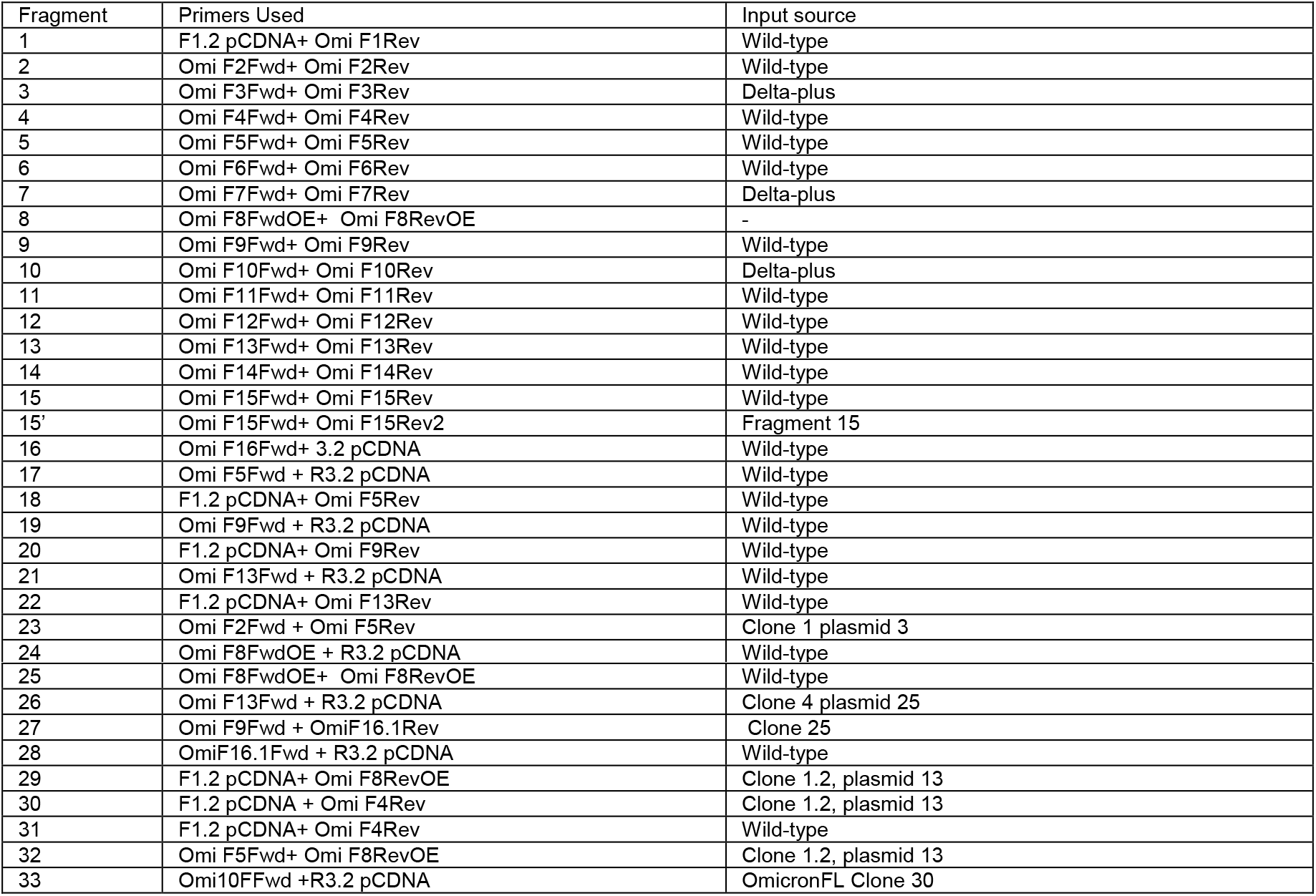
A list of fragments generated during the construction of the Omicron Spike clone, the primers and templates used for their generation

| Fragment | Primers Used | Input source |
| --- | --- | --- |
| 1 | F1.2 pCDNA+ Omi F1Rev | Wild-type |
| 2 | Omi F2Fwd+ Omi F2Rev | Wild-type |
| 3 | Omi F3Fwd+ Omi F3Rev | Delta-plus |
| 4 | Omi F4Fwd+ Omi F4Rev | Wild-type |
| 5 | Omi F5Fwd+ Omi F5Rev | Wild-type |
| 6 | Omi F6Fwd+ Omi F6Rev | Wild-type |
| 7 | Omi F7Fwd+ Omi F7Rev | Delta-plus |
| 8 | Omi F8FwdOE+ Omi F8RevOE | - |
| 9 | Omi F9Fwd+ Omi F9Rev | Wild-type |
| 10 | Omi F10Fwd+ Omi F10Rev | Delta-plus |
| 11 | Omi F11Fwd+ Omi F11Rev | Wild-type |
| 12 | Omi F12Fwd+ Omi F12Rev | Wild-type |
| 13 | Omi F13Fwd+ Omi F13Rev | Wild-type |
| 14 | Omi F14Fwd+ Omi F14Rev | Wild-type |
| 15 | Omi F15Fwd+ Omi F15Rev | Wild-type |
| 15' | Omi F15Fwd+ Omi F15Rev2 | Fragment 15 |
| 16 | Omi F16Fwd+ 3.2 pCDNA | Wild-type |
| 17 | Omi F5Fwd + R3.2 pCDNA | Wild-type |
| 18 | F1.2 pCDNA+ Omi F5Rev | Wild-type |
| 19 | Omi F9Fwd + R3.2 pCDNA | Wild-type |
| 20 | F1.2 pCDNA+ Omi F9Rev | Wild-type |
| 21 | Omi F13Fwd + R3.2 pCDNA | Wild-type |
| 22 | F1.2 pCDNA+ Omi F13Rev | Wild-type |
| 23 | Omi F2Fwd + Omi F5Rev | Clone 1 plasmid 3 |
| 24 | Omi F8FwdOE + R3.2 pCDNA | Wild-type |
| 25 | Omi F8FwdOE+ Omi F8RevOE | Wild-type |
| 26 | Omi F13Fwd + R3.2 pCDNA | Clone 4 plasmid 25 |
| 27 | Omi F9Fwd + OmiF16.1Rev | Clone 25 |
| 28 | OmiF16.1Fwd + R3.2 pCDNA | Wild-type |
| 29 | F1.2 pCDNA+ Omi F8RevOE | Clone 1.2, plasmid 13 |
| 30 | F1.2 pCDNA + Omi F4Rev | Clone 1.2, plasmid 13 |
| 31 | F1.2 pCDNA+ Omi F4Rev | Wild-type |
| 32 | Omi F5Fwd+ Omi F8RevOE | Clone 1.2, plasmid 13 |
| 33 | Omi10FFwd +R3.2 pCDNA | OmicronFL Clone 30 |

**Table 3:** Timeline for the synthesis of the Omicron Spike construct.

| *Day* | *Activity* |
| --- | --- |
| 1 | Strategy finalized |
| 2 | Primers ordered |
| 3 |  |
| 4 |  |
| 5 | Primers received |
| 6 | Fragment PCRs round 1 |
| 7 | OE and Gibson assembly round 1 |
| 8 | Col PCR 1 |
| 9 | Plasmid prep, sent for sequencing |
| 10 |  |
| 11 |  |
| 12 |  |
| 13 | Sequencing results; Fragment PCRs round 2 |
| 14 | Round 2 cloning |
| 15 |  |
| 16 | Col PCR 2 |
| 17 | Plasmid prep, sent for sequencing |
| 18 |  |
| 19 |  |
| 20 | Sequencing results |
| 21 | Fragment PCRs round 3 |
| 22 | Round 3 cloning |
| 23 | Col PCR 3 |
| 24 |  |
| 25 |  |
| 26 | Plasmid prep, sent for sequencing |
| 27 |  |
| 28 |  |
| 29 | Sequencing results for final clone |

## Overall strategy

The comprehensive list of mutations incorporated into the Omicron Spike were based on the GISAID sequence EPI_ISL_6814922 (2021-11-27)[4] and are listed below:

Spike A67V, Spike H69del, Spike V70del, Spike T95I, Spike G142D, Spike V143del, Spike Y144del, Spike Y145del, N211del, Spike L212I, Spike ins214EPE, Spike G339D, Spike S371L, Spike S373P, Spike S375F, Spike K417N, Spike N440K, Spike G446S, Spike S477N, Spike T478K, Spike E484A, Spike Q493R, Spike G496S, Spike Q498R, Spike N501Y, Spike Y505H, Spike T547K, Spike D614G, Spike H655Y, Spike N679K, Spike P681H, Spike N764K, Spike D796Y, Spike N856K, Spike Q954H, Spike N969K, Spike L981F

The overall strategy was to split the Spike-coding sequence into fragments, each to be amplified by PCR, incorporating the desired mutations in the primer binding regions. The fragments would then be stitched together by overlap PCR, and then assembled by Gibson assembly [2] to create the complete sequence (Figure 1). Gibson assembly uses a combination of three enzymes to ligate fragments with homologous ends, and leaves no scar in the final ligated product, an essential requirement while assembling a large number of fragments. The process efficiently ligates up to five fragments, after which the efficiency drops. Since this involved a large number of fragments, at least two rounds of assembly and verification by sequencing were anticipated.

**Figure 1:**
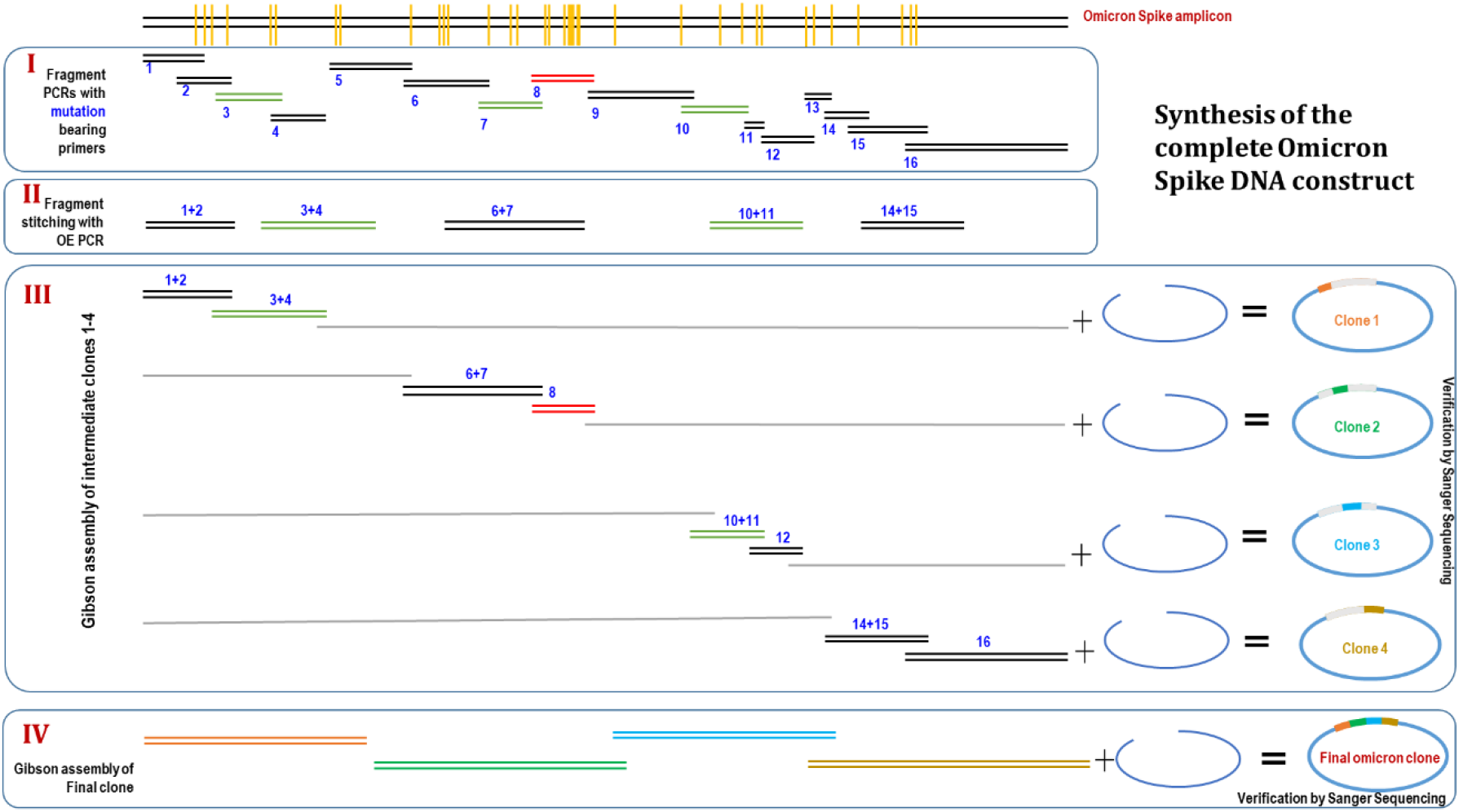
Strategy for the construction of Omicron Spike incorporating 37 mutations.

We used the Spike coding sequence, with the 19 amino acid deletion at the C-terminal end, as our template since this was the construct to be used in the Pseudovirus neutralization assay[5]. We used the wild-type Spike, or the Delta-plus variant Spike DNA as the template for PCR, based on the mutations required to be incorporated in a specific fragment. Three mutations were incorporated from the Delta-plus Spike DNA template and the rest were introduced using primers. We divided the Spike sequence into 16 fragments, each demarcated by the location of one or more Omicron specific mutations, and each overlapping with the preceding and following fragments by at least 27bp. We designed primers for amplification of each fragment incorporating the required mutation(s) at the center of the primer, with at least 12bp of perfectly matched sequence to promote proper annealing and amplification during PCR (Figure 1).

There were two regions that required a slightly different design. Fragment 8 contained nine mutations spread over ~200bp region, which did not align with the aforementioned strategy. Therefore, two long oligonucleotides with a 23bp overlap were designed to generate fragment 8 by routine overlap-extension. Fragment 15 contained two mutations spread over 48bp, which was too short to split into separate fragments, and too long for a single primer (~74bp). Hence, two consecutive reverse primers were designed to be used sequentially for fragment 15, and the second PCR was done using the first PCR as template.

A few additional modifications were employed to simplify the assembly and screening process:

1. Plasmids (pCMV-Spike (wt) and pCDNA-Spike (Delta-plus) used as a template for PCRs were digested with at least two restriction enzymes in order to reduce the background from inserts during transformation.
2. pCMV-Spike was used as the template for PCRs, and fragments were cloned into pCDNA3.1. This allowed for the correct identification of clones by colony PCR using a combination of Spike-specific internal primers and pCDNA3.1 specific primers at each end.
3. The last mutation L981F was incorporated only in the final round of assembly, so as to be able to amplify one end from a pCMV-Spike template and thus allowing colony PCR based screening as mentioned in #2 above.
4. The assembly was planned in such a way as to generate intermediate chimeric clones, which would contain the Omicron mutations in just the NTD, RBD or CTD (with the rest being wild-type Spike sequence).

## Methods and Results

The complete Omicron Spike DNA construct was split into four parts for ease of synthesis, based on the technical limitation of the Gibson assembly process. The mutation containing fragments of each part were amplified first, and assembled into a clone in the first round. A simple workflow for the generation of each clone is shown below (Figure 2). The four parts were then combined into the complete Omicron Spike DNA construct, by following the same workflow in subsequent rounds.

**Figure 2:**
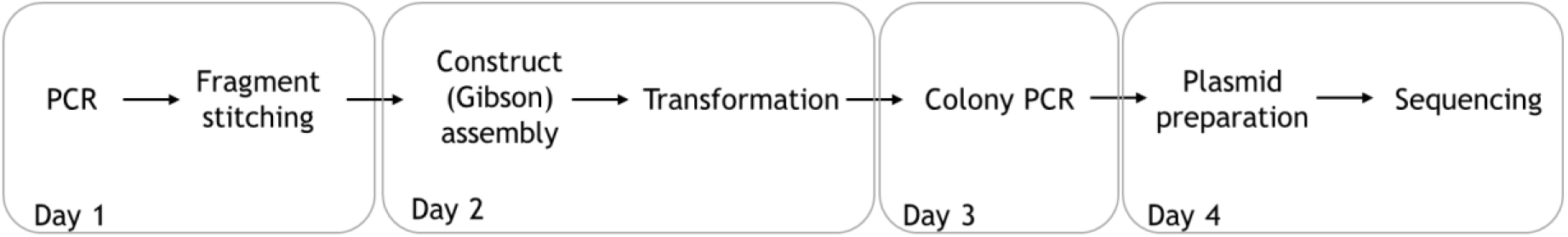
A simple workflow for the assembly of each part of the complete Omicron construct.

### PCR amplification of fragments

Each of the fragments (1-7, 9-16) was amplified from the appropriate template in a standard 25μl PCR reaction (Figure 3A). Taq polymerase was used for PCR amplification of fragments for the first part (fragments less than 1000bp). For later rounds, Q5 polymerase was used to avoid PCR – errors. Additional fragments to complete the assembly of each clone were also amplified using the pCMV-Spike (wild-type) template (labeled fragments 17-23). For example, clone 1 would include mutation containing fragments 1,2,3,4 and the rest (5-16) amplified from wild-type Spike (Fragment 17). The list of primers used in this study can be found in Table 1, and details of the fragments used, along with the primers and template DNA used to amplify them, can be found in Table 2. A standard PCR reaction was assembled as follows:

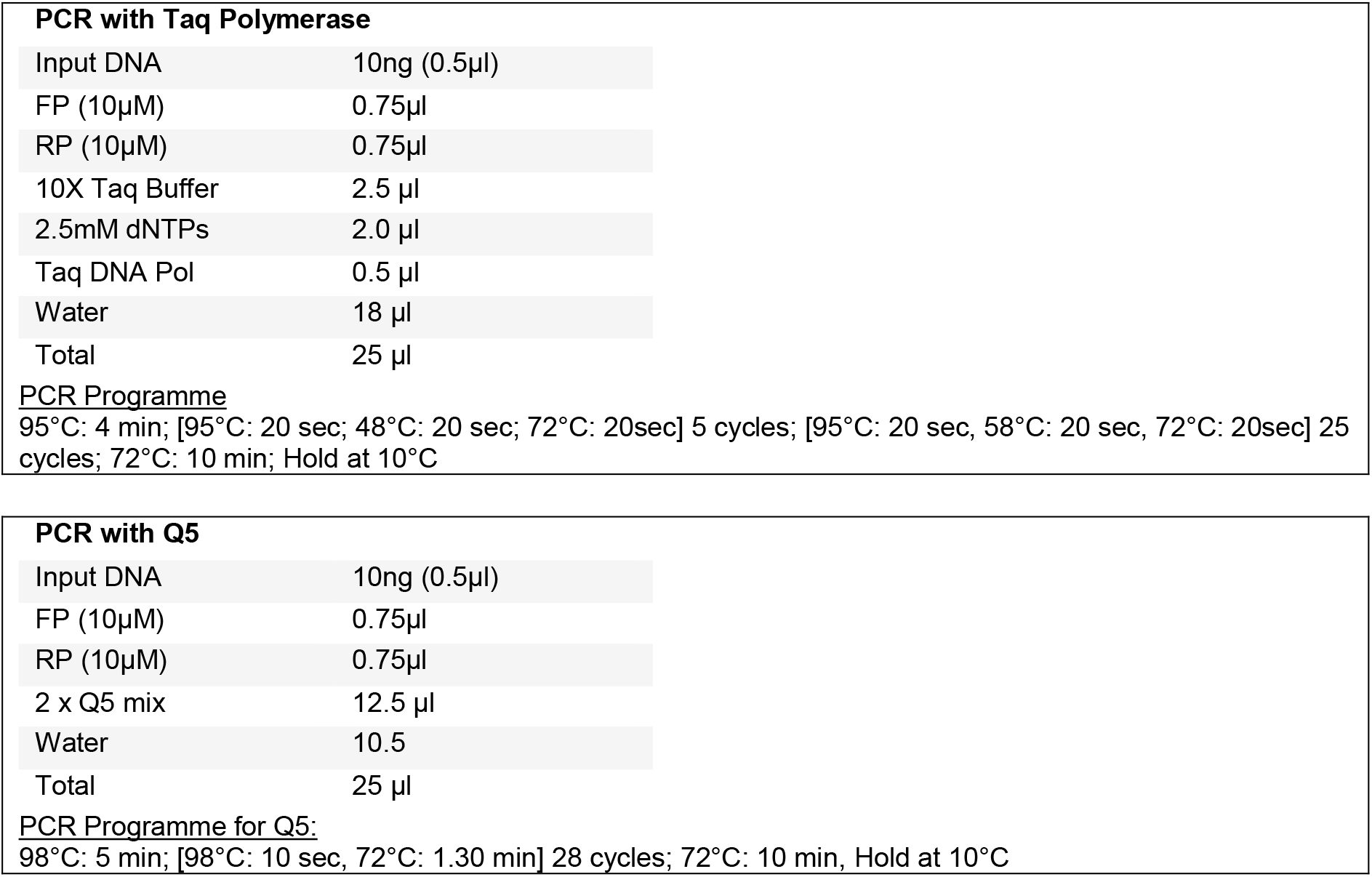

**Figure 3:**
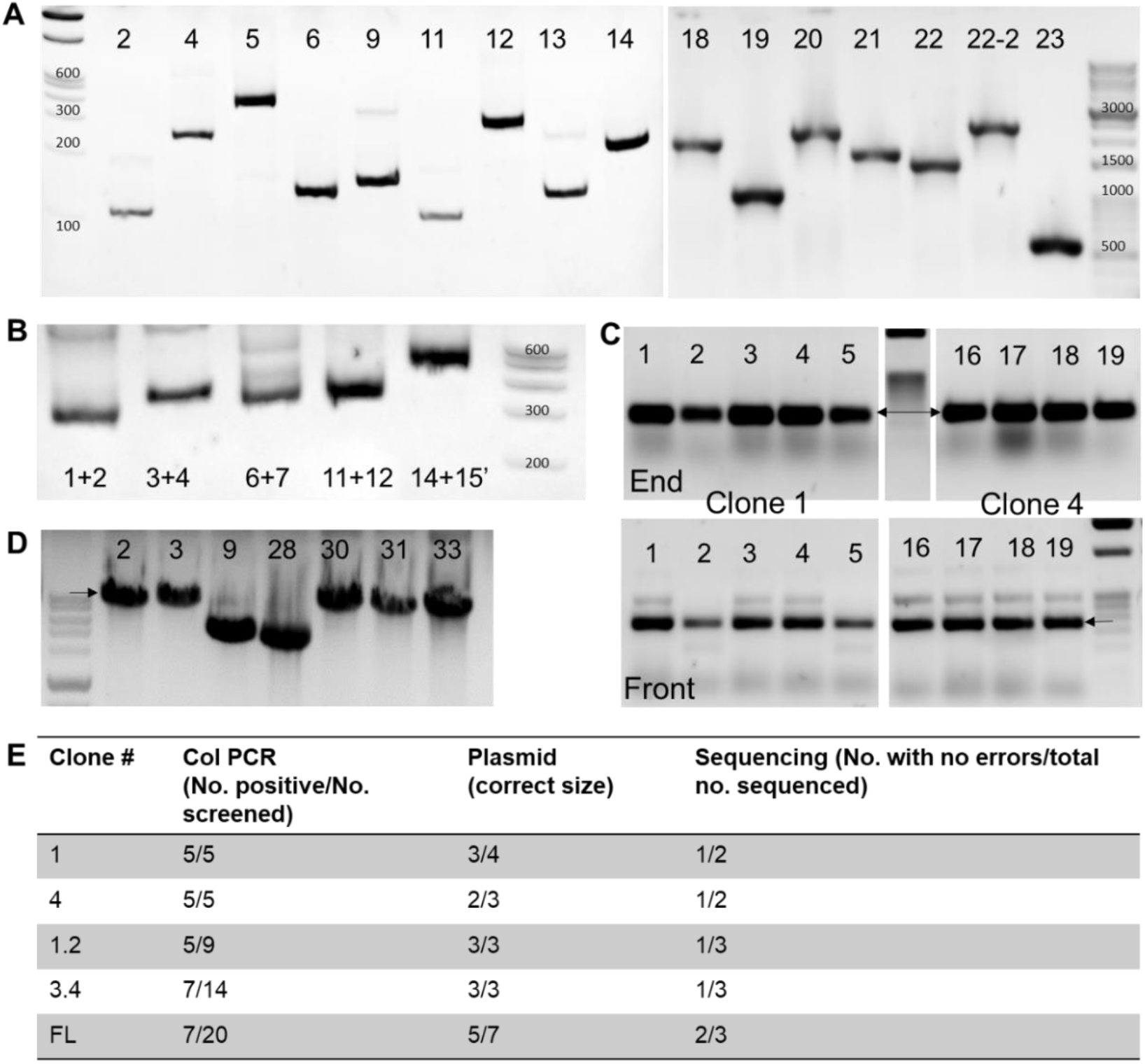
Representative gel images at various stages of Omicron synthesis. **A.** Individual fragments were amplified by PCR using Taq Polymerase or Q5 and 8 μl was visualised on a native polyacrylamide gel (fragments <800bp) or agarose gel. **B.** Fragments stitched together by OE were visualised by native PAGE. **C.** Colony PCR was used to confirm the presence of the complete insert using two primer sets, one across the vector-insert junction at the N terminal end of the Spike ORF (Front), and the second at the junction near the C terminal end (End). Two representative clones are shown. **D.** Plasmids amplified from positive colonies were checked for size to confirm the presence of all insert fragments, by resolving on an agarose gel. E. For each clone assembled using Gibson assembly, the fraction of positive colonies by colony PCR, by plasmid size and finally by sequencing are indicated as a measure of efficiency of the cloning process.

#### Challenge

The overlap PCR to generate fragment 8 was inefficient, and prevented efficient assembly into Clone 2. This was corrected in the subsequent round of assembly by using the long oligonucleotides as just primers.

### Fragment stitching

In order to improve the efficiency of Gibson assembly by reducing the number of inserts (to a maximum of 3 excluding vector), we stitched short PCR fragments using the method of Hilgarth et al [6]. Briefly, each PCR fragment was purified on a DNA cleanup column to get rid of primers, combined in equimolar amounts, and stitched together in overlap-extension 1 (OE1) 1. Subsequently the stitched fragment was amplified using the forward primer of the first fragment, and reverse primer of the second fragment in OE2 (Figure 3B).

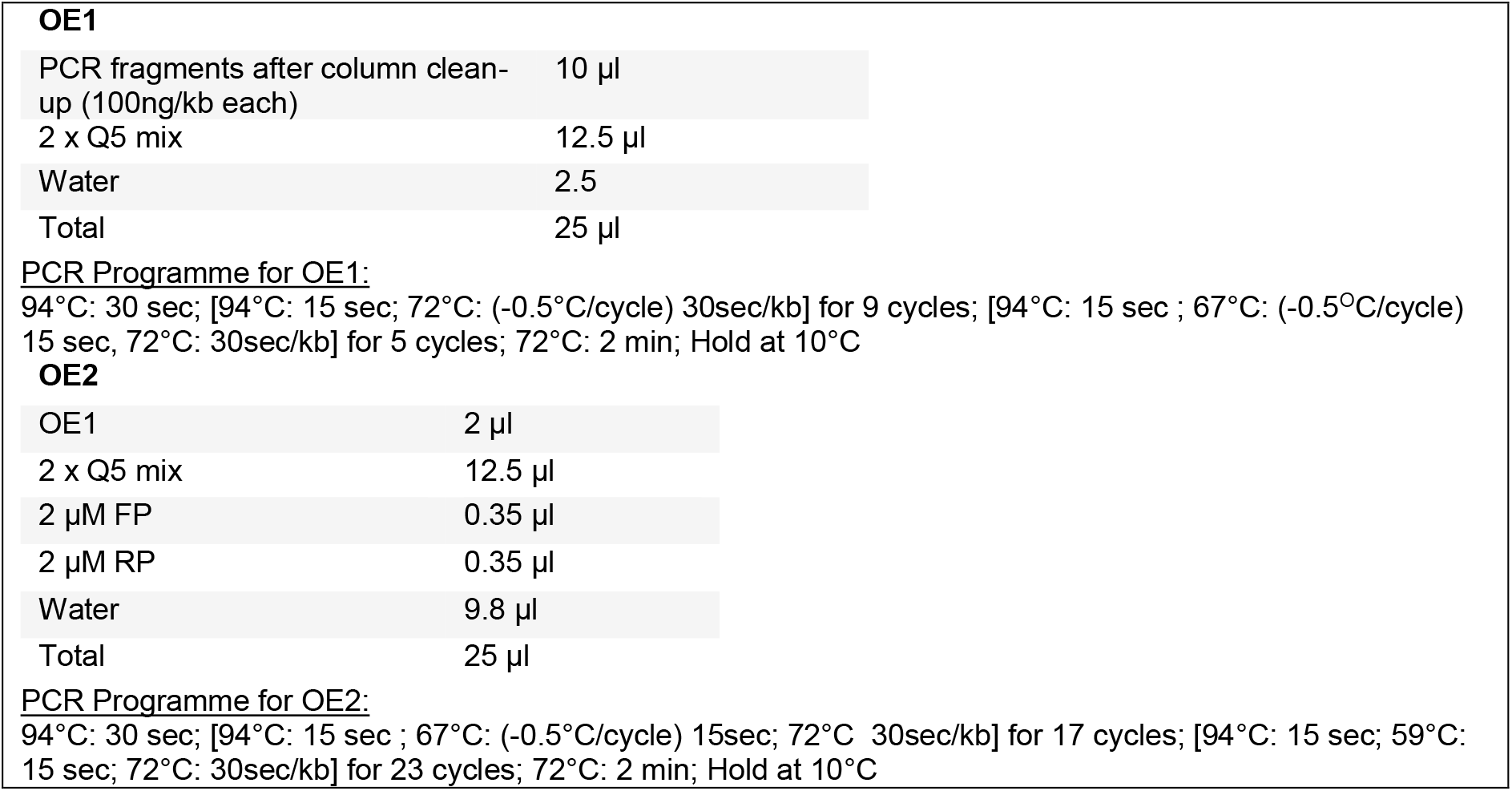

### Construct assembly Part 1- Gibson assembly of Clones 1,2,3 and 4

pCDNA3.1 was digested with BamHI and HindIII, treated with Antarctic Phosphatase and purified using a DNA cleanup column, and checked for background by transformation before using in the assembly reaction. Each of the clones was assembled with equimolar amounts of cut vector and inserts, except in the case of fragments shorter than 400bp, for which a 3-5 fold molar excess was used. The fragments that were incorporated into each of the clones are listed below and the details of the fragments can be found in Table 2. A typical Gibson assembly reaction was assembled as described below, incubated at 50°C for 1 hour, and transformed into 200μl competent cells. At least 40-50 colonies were obtained for each of the 4 reactions.

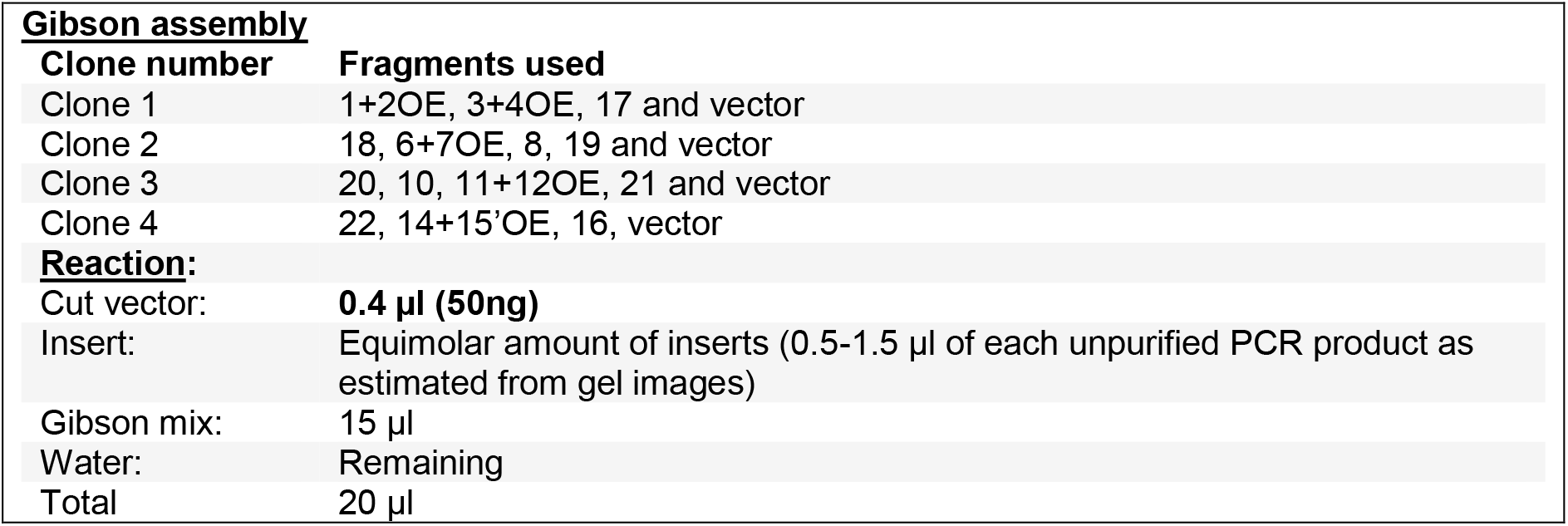

### Colony PCR screening

For each clone, 5 colonies were screened by colony PCR using two sets of primers: AS145 and AS645/ Omi F4Rev (Front), and AS575 and AS507 (End), to confirm insertion of the entire Spike sequence. Each colony was resuspended in 20 μl of water, lysed by heating, and 1.5 μl of the supernatant was used as input for a 10 μl PCR reaction. Plasmid was prepared from at least 2 positive clones, size-verified on a gel, and sent for sequencing with primers expected to cover the mutated region.

#### Challenge

Since Fragment 8 could not be generated efficiently, 10 colonies were screened for clone2, and 4 plasmids were sequenced. However, no mutations in fragment 8 were incorporated. In addition, for clone 3, all of the sequencing reactions failed. Therefore, we altered the strategy to include repeats of these two segments in the subsequent steps.

### Construct assembly Part 2- Gibson assembly to generate clone 1.2 (combining clones 1 and 2) and Clone 3.4 (clones 3 and 4)

For parts 2 and 3 of construct assembly, protocols identical to those described for part 1 above, were followed. The fragments assembled for each are listed below.

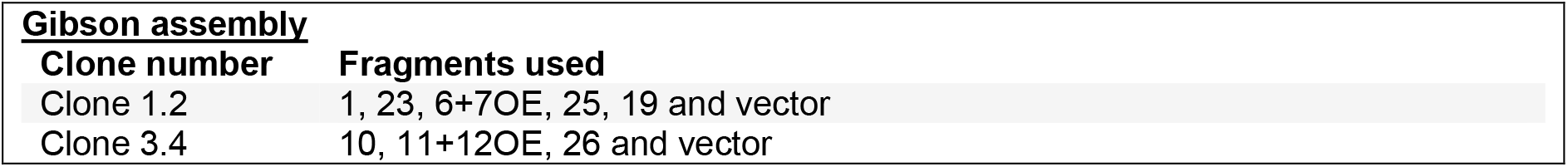

We obtained positive clones for both in this round, as confirmed by sequencing. A 6-fragment assembly (5 inserts and vector) proved to be successful as seen in the generation of clone 1.2.

### Construct assembly Part 3- Gibson assembly to generate full length clone

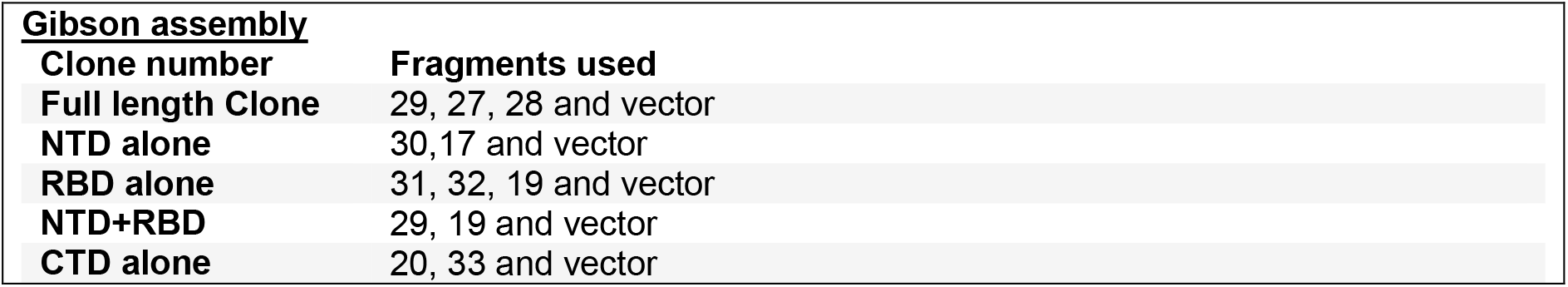

We obtained at least one positive clone for each of the above constructs, as confirmed by sequencing.

#### Challenges

In some instances, we observed errors in the primer binding regions which we attribute to errors in the oligonucleotides during synthesis, especially for long oligonucleotides (such as those for fragment 8). We overcame this by using a shorter oligonucleotide for a second round of PCR to eliminate specific errors.

## Discussion

In this study, we report a strategy for simultaneous site directed mutagenesis at multiple sites spread over several kilobases, with the added option of being able to generate a combinatorial library of mutant constructs. The method uses simple laboratory reagents and protocols and the strategy and methods are widely applicable. In addition to being an alternative for gene synthesis for complex constructs bearing multiple mutations, this technique may be used to synthesize a large collection of single as well as modular combinations of mutations. Constructs generated in this manner could be useful in a wide array of applications including but not limited to, (alanine) scanning mutagenesis for identification of sites of post-translation modifications [7], fitness, function, activity or structure contributions [8]–[10], as well as for functional analysis of a spectrum of patient mutations in genes implicated in monogenic disorders [11].

With the specific example of the Omicron Spike gene, we were able to generate the construct containing 37 mutations spread across a ~3.7kb DNA fragment using simple methods and reagents, accessible to most basic molecular biology labs in the world. Despite delays in sequencing turnaround, and strategic/experimental failures, we were able to complete this synthesis in half the time that it took to receive a commercially synthesized construct at our location. As suggested by the numbers in Figure 3E, the efficiency at each step was quite high and this complex construct with the desired sequence was assembled by screening only 5-10 colonies at each intermediate stage. Along the way, we were also able to generate constructs containing mutations only in smaller domains in the full-length gene.

A previous report described the LFEAP mutagenesis method which also relied on PCR and ligation mediated assembly to incorporate multiple mutations [12]. However, this method involved the amplification of the entire plasmid, bringing with it, the increased possibility of errors, and T4 DNA ligase mediated assembly which is usually less efficient than overlap-mediated assembly. Traditional scanning mutagenesis methods involve elegant but technically more complex strategies using special modified templates [13]. Gibson assembly was the single most important experimental method that made the current strategy possible. This method allows for seamless assembly without the requirement for or introduction of any extra or restricted nucleotide sequences. Efficient incorporation of up to five inserts in one assembly, significantly reduces the amount of time required to assemble complex constructs. Combined with the very efficient oligo stitching method reported by Hilgarth et al, technically up to ten fragments may be assembled in one shot.

In this study, we have demonstrated single or multiple amino acid changes at consecutive or spaced locations, insertions and deletions (single or up to 3 residues). We expect that large deletions (>10bp) will also be technically feasible. However, introduction of repetitive sequences, or large insertions may be slightly more challenging and may require modification of the described strategy. Despite these caveats, we believe that this strategy will be of great use in resource-constrained settings.

## Author contributions statement

R.R., A.S., K.P and K.C. conceptualized the study. R.R. and A.S. designed and executed the experiments, and analyzed the results. R.R and A.S. wrote the manuscript with inputs from K.P and K.C.

